# Mean growth rate when rare is not a reliable metric for persistence of species

**DOI:** 10.1101/762401

**Authors:** Jayant Pande, Tak Fung, Ryan Chisholm, Nadav M. Shnerb

## Abstract

The coexistence of many species within ecological communities poses a long-standing theoretical puzzle. Modern coexistence theory (MCT) and related techniques explore this phenomenon by examining the chance of a species population growing from rarity in the presence of all other species. The mean growth rate when rare, 𝔼[*r*], is used in MCT as a metric that measures persistence properties (like invasibility or time to extinction) of a population. Here we critique this reliance on 𝔼[*r*] and show that it fails to capture the effect of random abundance variations on persistence properties. The problem becomes particularly severe when an increase in the amplitude of stochastic temporal environmental variations leads to an increase in 𝔼[*r*], since at the same time it enhances random abundance fluctuations and the two effects are inherently intertwined. In this case, the chance of invasion and the mean extinction time of a population may even go down as 𝔼[*r*] increases.

## I. INTRODUCTION

Understanding the factors and processes that shape the diversity of species in ecological communities is among the most important questions in ecology. The prevalence of highly diverse communities, in which many species interact with each other and with abiotic factors, has long been a source of wonder as it seems to violate the competitive exclusion principle [1, 2] and/or May’s negative relationship between complexity and diversity [3]. Many possible mechanisms that may explain the maintenance of species diversity have been suggested [4], but the identification of those which actually influence a given system is still a formidable task.

Modern coexistence theory [4–9] (MCT) is a widely-used conceptual framework aimed at clarifying the conditions for species coexistence. Instead of fully considering the intricate network of dynamical interdependencies between all the species populations in a community, MCT simplifies the coexistence problem tremendously by focussing on the invasibility of a single species population, i.e., on the chance of a single invading population (or a small group of invading populations) to establish, given the dynamics of all the other resident species and the external environment. Invasibility is related to persistence time: if a population tends to recover from low frequencies, then its time to extinction is taken to be large [10]. Two populations coexist if each of them invades the other, and analogously, the persistence of high-diversity assemblages is examined by looking at the chance of each of the species to invade the community [11]. Importantly, MCT explicitly considers stochastic temporal environmental fluctuations, which have been found to be a key driver of community dynamics in natural communities [12–15].

In MCT, invasibility is measured by a single metric, 𝔼[*r*], which is the mean growth rate of a population when rare – i.e., when its relative abundance (frequency) is so low that it does not affect the dynamics of all the other populations. Given a time series of (low) frequencies {*x*_*t*_, *x*_*t*+Δ*t*_, *x*_*t*+2Δ*t*_…}, this mean growth rate is defined as [6]

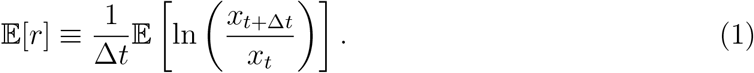

Here we would like to critique the reliance of MCT (and related theories) on the metric 𝔼[*r*] as a quantitative measure for persistence. We shall argue that, while the **sign** of 𝔼[*r*] is indeed an important characteristic of persistence, the **magnitude** of 𝔼[*r*] does not relate directly to persistence. Once 𝔼[*r*] > 0, a further increase in its value is not necessarily related to an increase in mean time to extinction or invasibility, and in some cases these quantities can even decrease when 𝔼[*r*] increases.

Accordingly, 𝔼[*r*] cannot be considered as a “general quantitative criterion for the persistence of a species” [5] and its value cannot be used to “decompose and compare” [8] the mechanisms that determine persistence and their relative importance. A contribution to 𝔼[*r*] is not necessarily a contribution to persistence, so one cannot quantify the contribution of a certain mechanism to persistence by comparing the value of 𝔼[*r*] in the presence and in the absence of this mechanism, which was the method used in [11] and in [16]. Similarly, 𝔼[*r*]-based metrics cannot be used for comparing the persistence properties of different communities, as done in [17].

The problem with the use of 𝔼[*r*] lies in its failure to reflect another important factor that affects persistence: the strength of random abundance fluctuations, or technically, the “diffusion” along the log-abundance axis. As we shall show below, when this diffusion is strong it may increasingly reduce the effect of 𝔼[*r*], making its value less and less relevant. Thus, invasibility and other persistence measures are associated with the relative strength of 𝔼[*r*] with respect to the random abundance variations.

The problem is highlighted when one considers the ability of temporal environmental stochasticity to promote coexistence, an effect that has attracted a lot of interest in the last few decades [4, 5, 8, 11, 18, 19]. In this case, stochastic fluctuations in the environment produce stochastic fluctuations in the per capita reproductive success of a species, and thereby govern both 𝔼[*r*] and the strength of the random abundance variations. This means that an increase in the amplitude of the stochastic environmental fluctuations increases both 𝔼[*r*] and the diffusion along the log-abundance axis, so that one cannot examine the two quantities separately.

Throughout this paper we consider the reliability of 𝔼[*r*] in different scenarios, mainly those in which stochastic environmental fluctuations promote coexistence. We begin with the Chesson-Warner lottery model [4, 5, 18] and examine it first for an infinite population and then for a finite population (i.e., with demographic stochasticity). We also study related models, like the forest dynamics model of [17, 19], and comment on related metrics, like Δ*I*_*b,i*_ used in [11]. In all these cases, we show that 𝔼[*r*] cannot be used as a single indicator for persistence properties.

## II. METHODS

The two main characteristics of stochastic temporal environmental fluctuations are their typical amplitude *σ* and typical duration (correlation time) *δ* [20]. If the fitness is related to the mean number of offspring (or seeds or larvae) per individual, this mean is taken to be 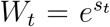, where *s*_*t*_ is a stochastic process with variance *σ*^2^ and correlation time *δ*. All times are measured in units of the generation time.

We examine two measures of persistence: the chance of invasion from rarity and the time to extinction of an already established population. The chance of invasion is the probability of a rare population to reach a given density before going extinct. This property depends only on the probabilities of transition between different states of the system, and is independent of the overall rate of events: for a given stochastic process, if all rates are doubled or halved, the chance of invasion remains the same. On the other hand, the time to extinction of an established population depends on both the transition probabilities and on their rates: when all rates are doubled, the time to extinction is halved.

To assess the use of 𝔼[*r*], we perform numerical experiments and examine the chance of invasion from rarity and the time to extinction of an established population when *σ* and *δ* vary across experiments. For 𝔼[*r*] to be used as a single metric for persistence, one should expect the following:

1. The persistence properties increase monotonically with 𝔼[*r*].
2. The persistence properties are the same for different combinations of *δ* and *σ*^2^, when these combinations yield the same 𝔼[*r*].

As we shall see, this is not the case. In the presence of stochastic environmental fluctuations the magnitude of 𝔼[*r*] tends to increase with *σ* and to decrease with *δ*, but the actual persistence properties may respond differently. Even when the response is in the “correct” direction, condition (2) above is not fulfilled.

When the amplitude of random abundance variations is small, one can implement the diffusion approximation [20, 21]. The theory of stochasticity-induced stabilisation is considered within the diffusion approximation framework in [22]. Here (in Supplementaries I and II) we reproduce some of these results and use them to obtain some insight into the failure of 𝔼[*r*], to suggest alternative metrics and to compare (when possible) our numerical results with analytical predictions. In particular, when the diffusion approximation is applicable (i.e., for small *σ*) the association mentioned in the Introduction above becomes a direct relationship: both the chance of invasion and the mean time to extinction are determined by 𝔼[*r*]/*g*, where *g* is the strength of the abundance variations.

Most of the parameter space surveyed in our numerical experiments lies outside the region in which the diffusion approximation holds. Nevertheless, the insights gained using the diffusion approximation (i.e., in the small-*σ* regime) provide a qualitative understanding of the general relationships between 𝔼[*r*], abundance variations and persistence.

## III. RESULTS

### A. The lottery model: Extinction time, invasibility and 𝔼[*r*] in a two-species community

Persistence is related to the chance of invasion (the probability that a rare population will grow and reach some threshold abundance or frequency) and to the time to extinction of an established population. Both these quantities have to do with the dynamics of a population close to its extinction state (in other words, in an “extinction zone”). In reality any population is a collection of *n* discrete individuals, and the frequency *x* = *n/N* reflects the fraction of a given population in an entire community of *N* individuals. In this case, the extinction zone is naturally defined as the state with one individual, or only a few individuals. For theories that do not take into account the discreteness of individuals (i.e. theories without demographic stochasticity), the definition of the extinction zone is arbitrary, say 0 ≤ x ≤ ϵ where *E* is some small number, but to make sense in practical situations one should posit this *ϵ* to be of order 1/*N*.

What are the relationships between the metric 𝔼[*r*] and persistence? To examine this, let us look at the most famous example of stochasticity-induced stabilisation, the two-species Chesson-Warner lottery model [18]. In a single time-step (without loss of generality, we use yearly time-steps) the frequency of the focal species *x* satisfies

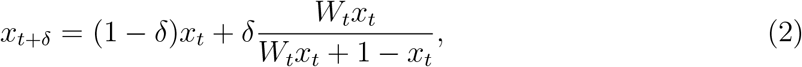

where *t* is time measured in units of the generation time, *δ* is the fraction of the individuals in the community dying in each year and *W*_*t*_ is the relative fitness of the focal species with respect to its rival species at time *t* (reflecting, for example, the mean number of seeds produced by a single individual of the focal species per one seed produced by the rival species). Under these dynamics the lifetime of adults is distributed geometrically with mean 1/*δ* years, so 1/*δ* is the generation time in years. If *W*_*t*_ is picked at random in each year, then *δ* is the correlation time of the environment as measured in units of the generation time.

When the focal species is rare (*x*_*t*_ ≪ 1, 1 − *x*_*t*_ ≈ 1),

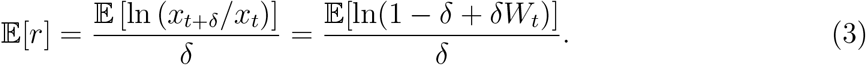

Clearly, the value of 𝔼[*r*] is determined by *δ* and by the statistical properties of *W*_*t*_. In what follows we consider the symmetric case, where the mean (over time) fitness of both species is the same and hence 𝔼[ln *W*_*t*_] = 0. In this case, we manipulate the amplitude of log-fitness variations *σ*^2^ = var (ln *W*_*t*_). The asymmetric case 𝔼[ln *W*_*t*_] ≠ 0 is considered in the Discussion section.

Fig. 1 shows how the invasibility and the time to absorption (mean time to fixation or extinction of the focal species, i.e., the mean time to extinction of either species), as measured in numerical simulations of the lottery process (2), are related to 𝔼[*r*]. Evidently, neither of the reliability conditions (presented in the Methods section above) is satisfied. Different combinations of *δ* and *σ*^2^ yield different persistence properties even if they correspond to the same 𝔼[*r*]. Moreover, not only does invasibility not grow monotonically with 𝔼[*r*], but the time to absorption even *decreases monotonically* as 𝔼[*r*] increases when *σ* is manipulated.

**FIG. 1:**
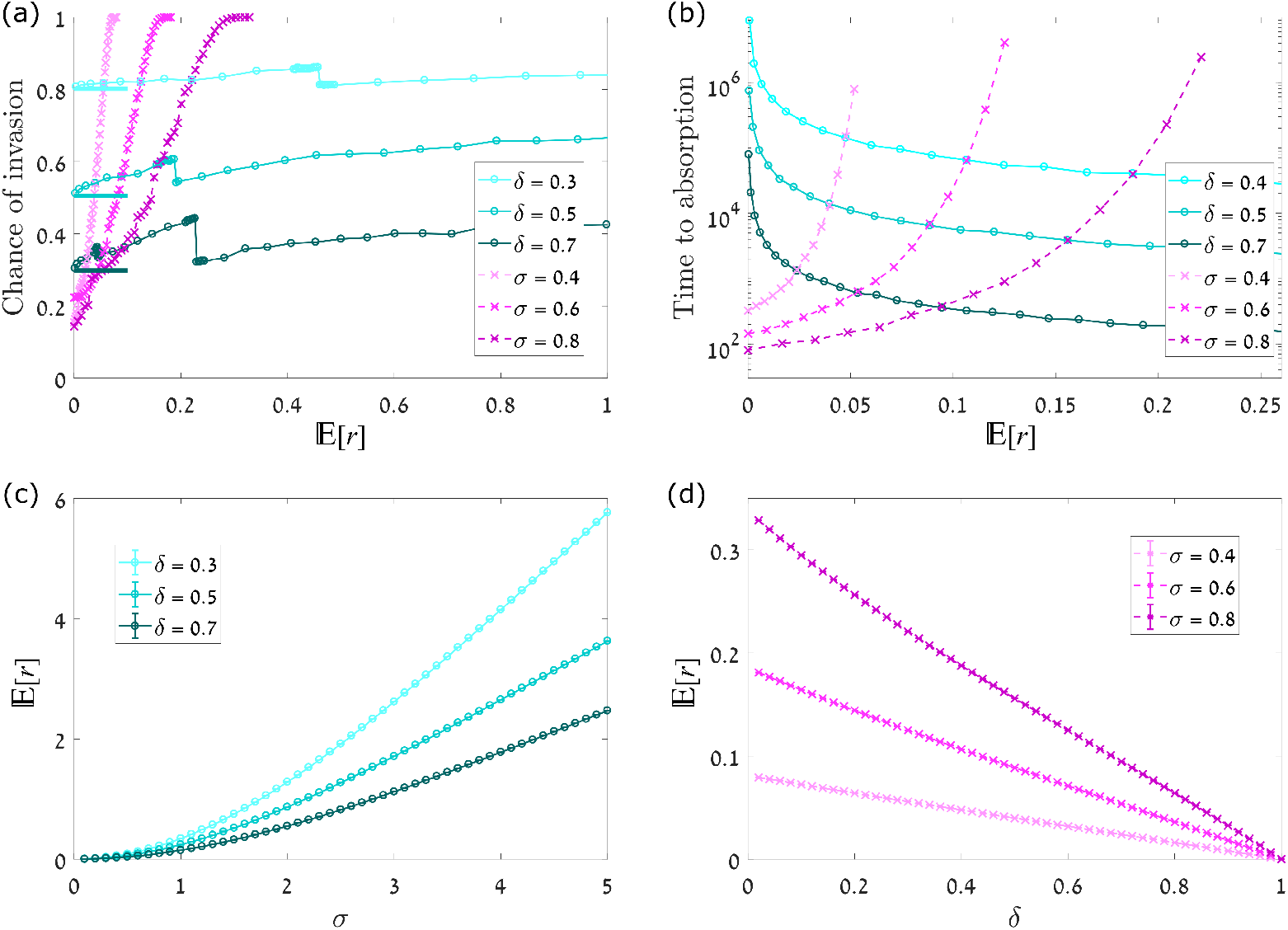
Persistence properties and 𝔼[*r*] for the symmetric two-species lottery model without demographic stochasticity, Eq. (2). The upper panels show the chance of invasion [panel (a)] and mean time to absorption [panel (b)] for different values of 𝔼[*r*]. The lower panels, (c) and (d), show the dependence of 𝔼[*r*] on the parameters *σ* and *δ* (amplitude and temporal correlation of stochastic environmental fluctuations, respectively). Circles (connected by full lines) represent the case where *δ* is held fixed and *σ* is varied, while crosses (connected by dashed lines) correspond to the reverse case. Clearly, different combinations of *σ* and *δ* which correspond to the same value of 𝔼[*r*] have completely different persistence properties. Moreover, when 𝔼[*r*] increases via an increase in *σ*, the chance of invasion is non-monotonic and the time to absorption decreases. Some features of these graphs are explained qualitatively, and even predicted quantitatively, by the diffusion approximation-based theory reviewed in Supplementary I. For example, the chance of invasion when *σ →* 0 fits perfectly the predictions of Eq. (6), marked by short horizontal strokes for different values of *δ* in panel (a). The jumps in the invasion curves are not numerical artefacts but may be traced to the chance of extinction in a single step or a few steps, see Supplementary I B. To find the chance of invasion, the process given by Eq. (2) was incremented from the initial condition *x* = 2*ϵ*, where *ϵ* = 0.001, with invasion being defined as reaching the state *x* = 0.1 before *x* = *ϵ*. To obtain the time to absorption, the same process was iterated from *x* = 0.5 until either *x* < *ϵ* or 1 *− x < E*. For both kinds of experiments, the fluctuations in the relative fitness of the focal species were assumed to be dichotomous (in Eq. (2), *W*_*t*_ = *e^±σ^*), and all results were averaged over 1000 simulation runs for each set of parameters.

#### Insights from the diffusion approximation

While the trends in Fig. 1 speak for themselves, one may understand them better by looking at the case where *σ*^2^ ≪ 1, i.e., when the diffusion approximation holds [21]. This case is analysed in detail in [22], and in Supplementary I we reproduce some of this analysis.

In this parameter regime the mean growth rate as defined in Eq. (3) is

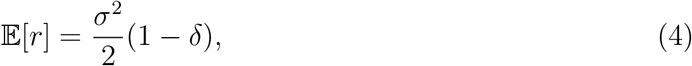

and the strength of stochastic fluctuations along the log-abundance axis, *g*, is equal to

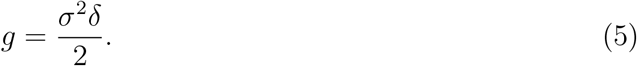

In Supplementary I [see derivation of Eq. (S9)] we show that in the diffusive regime the chance of invasion *ε*_+_ is given by

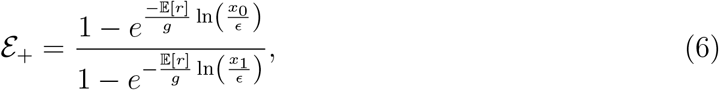

where *x* = *x*_0_ is the starting frequency, *x* = *ϵ* is the frequency at which extinction is defined, and *x* = *x*_1_ is the frequency at which invasion is defined to have occurred. Therefore, the chance of invasion is determined by the *ratio* 𝔼[*r*]/*g*, not by 𝔼[*r*] alone. The validity of Eq. (6) in the small-*σ* regime is demonstrated in Fig. 1(a). Since 𝔼[*r*]/*g* = (1 − *δ*)/*δ*, in this weak-stochasticity regime the chance of invasion is independent of *σ*. In contrast, 𝔼[*r*] grows like *σ*^2^, so its value may change without any corresponding change in *ε*_+_.

The rate of extinction events is proportional to the chance to find the system in the extinction zone [23]. When a stochastic system like (2) admits a coexistence state, it relaxes in the long run to an equilibrium state *P*_eq_(*x*) and forgets its initial condition. In Supplementary I we show that when *σ*^2^ *≪* 1 the chance to find the system in the extinction zone after relaxation is

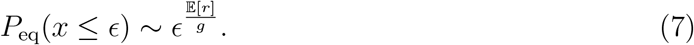

Again, the crucial quantity is the ratio 𝔼[*r*]/*g*, not 𝔼[*r*] itself, and when *σ* is small this chance is independent of *σ*.

The decrease in the time to absorption as *σ* increases, demonstrated in Fig. 1(b), is related to the overall rate of events which determines the relaxation times. The dynamics defined in Eq. (2) halt when *σ* = 0, so the time to equilibration diverges like 1/*σ*^2^ as σ → 0. The sign of 𝔼[*r*] is indeed an important qualitative feature of the system. As long as 𝔼[*r*] is positive, the chance of invasion *ε*_+_ is finite even when one invader is introduced in an infinite community, and the mean time to extinction of an established population grows with the size of the community *N* and diverges if this size goes to infinity. On the other hand, if 𝔼[*r*] is negative, a single invader cannot establish in an infinite community. This follows from Eq. (6): if *ε* is taken to be 1/*N*, the term 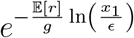 in Eq. (6) vanishes if 𝔼[*r*] is positive and diverges if 𝔼[*r*] is negative. Moreover, as explained in [22], the divergence of *P*_eq_(*x*) in the extinction zone [see Eq. (S6)] implies that the mean time to extinction becomes independent of the community size.

### B. The effect of demographic stochasticity

The lottery model (2) assumes that the frequency *x* may take any value. In practice, a community is made of an integer number *N* of individuals and *x* (which now equals *n/N*, where *n* is the number of individuals of the focal species) can change only in units of 1/*N*. The endogenous random birth-death process associated with this discreteness is known as demographic stochasticity (or ecological drift), and provides an extra source of abundance fluctuations, even if the environment is kept fixed [20].

In the absence of demographic stochasticity the lottery model (2) has two drawbacks: it does not have a naturally defined state of extinction (the frequency never reaches zero, so one has to impose a cutoff at *x* = *ϵ* as explained above), and its dynamics halt (extinction time diverges) when *σ* = 0. To consider a more realistic scenario we would like to examine the efficiency of 𝔼[*r*] as a measure of persistence when the lottery dynamics admit demographic stochasticity. When individuals are discrete the setup of an invasion experiment becomes natural: the extinction state is *n* = 0, and we measure the chance of a species with an initial population of a single individual to reach a finite fraction *x*_1_ = *n*_1_/*N* before extinction. The detailed definition of the model and the simulation procedure are given in Supplementary II.

A few typical results are depicted in Fig. 2. Clearly, the addition of demographic stochasticity does not fundamentally improve the usefulness of 𝔼[*r*] as an indicator of persistence. Again, the chance of invasion and the time to absorption are not unique functions of 𝔼[*r*], and an increase in 𝔼[*r*] may correspond to a decrease in these quantities, although the range of parameters for which this behaviour is observed is narrower than in the absence of demographic stochasticity.

**FIG. 2:**
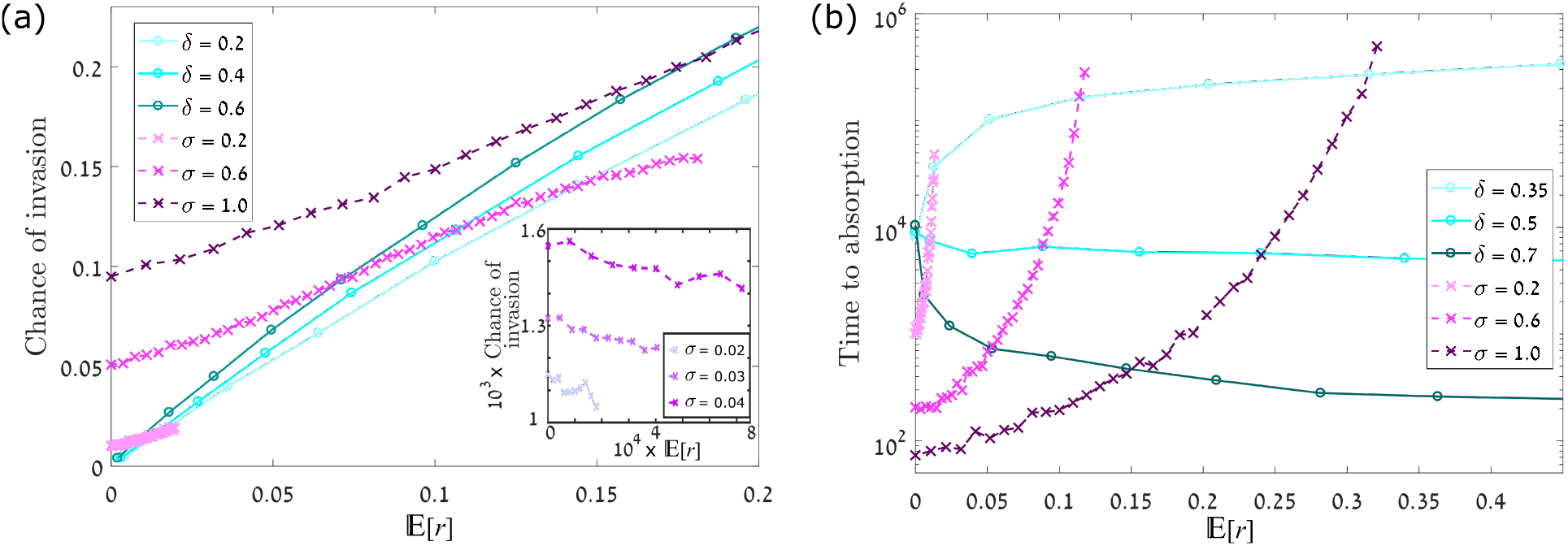
The chance of invasion [panel (a)] and the mean time to absorption [panel (b)] vs. 𝔼[*r*] as obtained from simulations of the lottery model with demographic stochasticity and *N* = 10000, for different values of *σ* and *δ* like in Fig. 1. The inset in panel (a) shows the chance of invasion for small values of *σ* as *δ* varies. In both the panels circles connected with full lines denote the case when *δ* is held fixed and *σ* is varied, while crosses connected with dashed lines denote the reverse case. The details of the simulation procedure are explained in Supplementary II. In invasion experiments the simulation is initiated at *n* = 1 and the chance to reach 0.1 *N* before extinction is plotted vs. 𝔼[*r*], the results averaged over 1000 simulation runs for each parameter set. For the time to absorption (extinction or fixation) experiments, the results are averaged for each parameter set over 100 different realisations starting from *n* = *N/*2, where now extinction and fixation are defined as the states *n* = 0 and *n* = *N*, respectively. As in the model without demographic stochasticity, different combinations of *σ* and *δ* yield different persistence properties even if they correspond to the same 𝔼[*r*], and an increase in 𝔼[*r*] does not necessarily imply a higher chance of invasion or longer absorption times. Moreover, while for most of the parameter space invasion becomes more likely as 𝔼[*r*] increases, for small enough values of *σ* the chance of invasion decreases with 𝔼[*r*] [inset, panel (a)].

#### Diffusion approximation insights

To obtain qualitative insight one may again use the diffusion approximation, as explained in Supplementary II. Demographic stochasticity provides an additional source of random abundance variations: in every time-step of the lottery dynamics, *δN* individuals are killed, so if *σ* = 0 the amplitude of abundance variations per time-step is 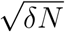. Accordingly, the variance of frequency fluctuations per time-step (of length *δ* generations) is *δ/N*, and the variance per unit time is 1/*N*. This extra term has to be included in the corresponding Fokker-Planck equation, see Supplementary II.

When *σ* = 0 (neutral dynamics), the only source of abundance variations is these demographic fluctuations. In this neutral limit they dictate the absorption time (which is of course independent of *δ*) and the chance of invasion [which is very small, 1/(*Nx*_*1*_)]. As a result, the mean times to absorption that were obtained from numerical experiments with fixed *δ* and varying *σ* (Fig. 2(b), circles joined by full lines) meet at the point *σ* = 𝔼[*r*] = 0, and the chance of invasion (Fig. 2(a), circles joined by full lines) approaches zero in this limit.

Even if *σ* is larger than zero, the strength of the random abundance variations is bounded from below by 1/*N* (see Eq. (S14) of Supplementary II). Accordingly, at very small values of *σ* (and *g*), Eq. (S14) suggests that the chance of invasion increases with *δ* (and decreases with 𝔼[*r*]), as demonstrated in the inset of Fig. 2(a). On the other hand, for larger values of *σ* our numerical experiments suggest that the chance of invasion decreases with *δ* (i.e., increases with 𝔼[*r*]). More work is needed to better characterise the relationship between the chance of invasion and 𝔼[*r*] in general scenarios (when the diffusion approximation does not hold, or when the system admits long-term memory as in the case of [17, 19] discussed below). However, the existence of even a small region where the chance of invasion runs counter to 𝔼[*r*] demonstrates the fallacy in relying on the latter as a measure of the former. With regard to the time to absorption (i.e., the time to extinction of either of the two species), an increase in *σ* has two opposite effects. On the one hand, it leads to an increase in the abundance variations, thus impeding coexistence and shortening absorption times. On the other hand, it increases the relative importance of stabilising mechanisms with respect to demographic stochasticity (ecological drift). As seen in Fig. 2(b), the net effect on the absorption time depends on the value of *δ* – with increasing *σ* (and 𝔼[*r*]), the absorption time increases when *δ* is low and decreases when *δ* is high.

Despite the ubiquity of demographic fluctuations, many studies focus on models without demographic stochasticity (like Eq. (2) above). The justification for this is that demographic stochasticity may be neglected if the community size *N* is large. Importantly, this argument does not hold for invasion experiments, since the system is initiated with a single individual, so the initial frequency scales with 1/*N*. As a result, our invasion curves converge to an *N*-independent limit when *N* → *∞* (as was checked numerically, results not shown), but this limit does not coincide with the curves of Fig. 1 which were obtained from Eq. (2) without demographic stochasticity.

### C. Related metrics for persistence

One of the main goals of MCT is to untangle the relative contribution of certain mechanisms to coexistence. To that aim, Ellner and coworkers [11] considered the role of the storage effect, i.e., the effect of the covariance between the environment and the competition strength (EC covariance), by comparing the value of 𝔼[*r*] in two numerical experiments, one with and one without EC covariance. This work is reviewed in Supplementary III. First, Ellner et al. simulated the dynamics of a two-species lottery model with correlations between the fitnesses of the two species (without demographic stochasticity), corresponding to the case with EC covariance, and calculated the value of 𝔼[*r*] in this case. Second, they removed the EC covariance and found the mean growth rate when rare in this case, denoted now by E[*r*^#^]. These authors have suggested the metric (see Supplementary III for technical details)

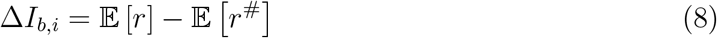

as a quantitative measure of the contribution of EC covariance to the persistence of a focal species, labelled with the subscript *i* (and with the subscript *b* standing for “between-species”).

Given the results presented above, it is clear that this 𝔼[*r*]-based metric Δ*I*_*b,i*_ does not reflect correctly the contribution of EC covariance to persistence. An increase in the contribution of EC covariance to the mean growth rate when rare does not simply translate to an increase in the actual persistence of a population. To decompose the effect of different mechanisms on persistence one would like to develop a similar procedure in order to assess, for example, the contribution of EC covariance to a metric which is more directly related to persistence, such as 𝔼[*r*]/*g* in the regime where *σ*^2^ is small enough.

Usinowicz et al. [17, 19] have examined the stabilising effect of stochastic environmental fluctuations by “neutralising” all other aspects of the community dynamics. To do that, they studied the dynamics of a tree community in which the mean growth, death and competition terms of all the tree species are exactly equal and the different species differ only in their (empirically calibrated) yearly fluctuations in seed, seedling or sapling production.

The persistence observed in a numerical experiment must be attributed to environmental fluctuations, since there is no other stabilising factor. The aim of [17] was to compare the effect of environmental fluctuations between different forests along the latitudinal gradient, so (unlike [11]) they did not try to disentangle the effect of different stabilising factors. The metric they used is *A*_*ij*_*A*_*ji*_ (reviewed in Supplementary IV). This metric is defined for a focal species *i* in competition with another species *j*, and is related to the *inverse* of the mean growth rate when the focal species is rare. Thus, when the value of *A*_*ij*_*A*_*ji*_ **decreases**, the absorption time and invasibility are supposed to increase.

To examine the use of this metric, we implemented the same forest dynamics model suggested in [17, 19]. We simulated a two species model, where stochastic fluctuations in the reproductive success are specified by the observed time-series of recruitment rates, *R*_*i*_(*t*), for two tree species (*Spondias mombin* and *Spondias radlkoferi*) over 14 years, as reported in [19]. We incorporated this dataset in a model with finite community size, *N* = 10000. As explained in Supplementary IV, we used a parameter *κ* to manipulate the amplitude of the environmental fluctuations (with *κ* analogous, but not identical, to *σ* above), such that *κ* = 1 gives environmental fluctuations which are the same as those reported in [19] whereas *κ* = 0 is the case with no environmental fluctuations. As *κ* varies, the amplitude of the environmental fluctuations changes, but all the other statistical properties of the environment (e.g. correlations) are left intact.

In Fig. 3, both *A*_*ij*_*A*_*ji*_ and the metrics that more directly measure persistence – namely, the chance of invasion and the mean absorption time – are plotted against *κ*. Again, there are no trivial relationships between the quantities. In particular, while *A*_*ij*_*A*_*ji*_ decreases monotonically with *κ* (indicating greater persistence), the mean absorption time may decrease. The chance of invasion does grow when *A*_*ij*_*A*_*ji*_ decreases, at least in the regime of parameters we have checked.

**FIG. 3:**
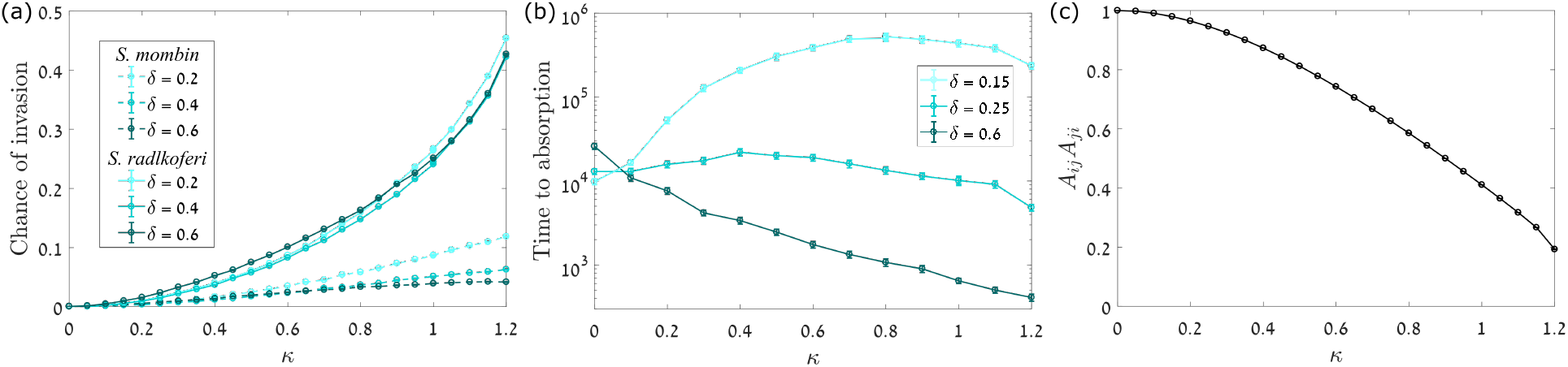
The chance of invasion [panel (a), dashed lines for *Spondias mombin*, full lines for *Spondias radlkoferi*] and the mean time to absorption [panel(b)] are plotted against the amplitude of stochastic environmental fluctuations *κ* for the forest dynamics model of [17, 19] for various values of *δ* (temporal correlation of environmental fluctuations). The case *κ* = 1 corresponds to the environmental stochasticity in [19] and we explore the parameter space for lower and higher stochasticity in the environment. The details of the model (which includes demographic stochasticity) and the simulation procedure are given in Supplementary IV. Note that for *κ ≥* 1.171 some of the modified recruitment rates [Eq. (S23)] become negative, so we restrict the plots to smaller *κ*-values. The dependence of the *A*_*ij*_*A*_*ji*_ metric used in [17, 19] on *κ* is shown in panel (c). The metric *A*_*ij*_*A*_*ji*_ decreases with increasing *κ* [panel (c)], and thus greater *κ* should indicate greater persistence properties. Instead, the mean time to absorption sometimes increases and sometimes decreases with *κ*. In this case, the chance of invasion indeed increases when *A*_*ij*_*A*_*ji*_ decreases.

## IV. DISCUSSION

Throughout this paper we have examined the utility of the mean growth rate of a species population when rare, 𝔼[*r*], as an indicator of the invasibility and the extinction time of the population in a community context. We suggest that the use of the sign of 𝔼[*r*] to identify a qualitative shift in persistence is correct, but this does not imply that the value of 𝔼[*r*] provides quantitative information about persistence. When the **sign** of 𝔼[*r*] is positive, the chance to find the population in the extinction zone goes to zero when the width of this zone goes to zero [24, 25]. If the extinction zone is defined as the region 0 ≤ x ≤ 1/*N*, it becomes narrower and narrower when the size of the community *N* increases. As a result, the mean time to extinction diverges with *N* for an established population. On the other hand, when 𝔼[*r*] is negative the chance to find the population in the extinction zone is finite even when the width of the extinction zone goes to zero, as the dynamics tend to take the population closer and closer to the limit of vanishingly small frequencies. In this case one expects an *N* - independent extinction time, as discussed in detail in [22]. This distinction between positive and negative 𝔼[*r*] was demonstrated explicitly in many studies of stochastic-logistic (and logistic-like) systems [20, 26–28].

However, while the sign of 𝔼[*r*] provides a fair binary classification scheme, the assumption that higher values of 𝔼[*r*] imply greater persistence (as measured by, for example, the chance of a population visiting the extinction zone or invading a community) is very problematic.

As a simple example one may think about stochastic-logistic systems [20, 26–28]. In these systems the intrinsic growth rate of the focal species, *r*(*t*), fluctuates in time, such that its mean is 𝔼[*r*], its variance is *σ*^2^ and the typical correlation time of the environment is *δ*. The mean time to extinction when *N* is large and 𝔼[*r*] > 0 scales like *N*^𝔼[*r*]/*g*^, where, as above, *g* = *σ*^2^*δ*/2 measures the strength of the stochastic abundance fluctuations. Accordingly, two systems with the same 𝔼[*r*] may have different persistence properties if the value of *g* is different.

For stochastic-logistic dynamics it is still true that an increase in 𝔼[*r*] leads to an increase in the persistence *if g* is kept fixed. But for dynamics whereby stochastic environmental fluctuations produce a stabilising mechanism that promotes coexistence of species, the situation is even more intricate: parameters like *δ* and *σ* control both the amplitude of the stochastic abundance variations and (through the stochasticity-induced stabilising mechanism, like the storage effect) the value of 𝔼[*r*]. As the same environmental factors govern the mean growth rate and the amplitude of the abundance fluctuations, one cannot consider these quantities independently anymore. As shown throughout this paper, this implies that 𝔼[*r*] is not a reliable metric for persistence.

Above we considered only communities with symmetric competition (i.e., with both species having the same mean fitness, or *s*_0_ ≡ 𝔼[ln *W*_*t*_] = 0). We have verified that for non-zero values of *s*_0_ it is still true that the persistence may be a non-monotonic function of 𝔼[*r*] (Fig. 4).

**FIG. 4:**
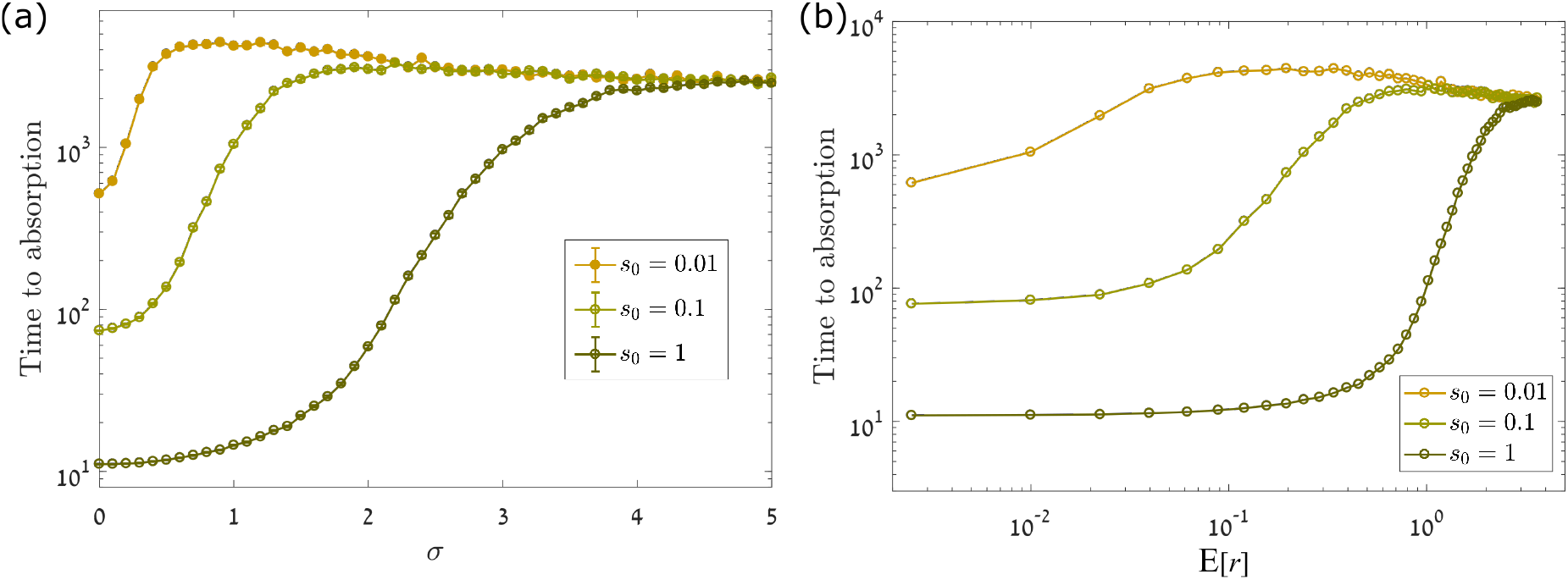
The time to absorption (extinction or fixation) for an asymmetric two-species lottery model with demographic stochasticity, with *N* = 10000 and *δ* = 0.5, plotted against (a) the parameter *σ* (amplitude of environmental fluctuations), and (b) the metric 𝔼[*r*]. The asymmetry in the model arises from the fitness advantage of the focal species over the other species, with the mean fitness advantage given by *s*_0_ *≡* 𝔼[ln *W*_*t*_] / = 0. As for the symmetric model in Fig. 2, the starting population of the focal species (and also of the competing species) is *n* = *N/*2, and extinction and fixation are defined to be the states *n* = 0 and *n* = *N*, respectively. The time to absorption, as in the symmetric case, can rise or fall with 𝔼[*r*] and *σ*.

In this study we considered only two-species communities in order to clearly highlight the problems with using 𝔼[*r*] to measure persistence. Although we did not simulate here the dynamics of a community with many species, we note that there are some recently-published simulation results (Figs. 5 and 6 of [29]) which show the species richness in high-diversity communities as a function of *σ*^2^. The model considered in [29] supports the storage effect which leads to stochasticity-induced stabilisation, so the mean growth rate when rare (for each population) increases with the amplitude of environmental variations *σ*. However the species richness decreases with *σ* if the correlation time *δ* is long.

Is there any alternative indicator to 𝔼[*r*]? As we have seen, when the diffusion approximation holds, the parameter 𝔼[*r*]/*g* governs the chance of invasion and the mean time to extinction. A recent work [28] suggests that when the dynamics of a population (when rare) are logistic or logistic-like, the persistence properties are determined by the parameter *q* which satisfies the equation

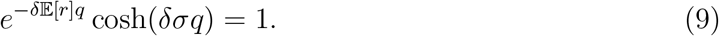

The value of *q* converges to 𝔼[*r*]/*g* when the diffusion approximation holds, and the time to extinction scales like *N* ^*q*^ even when the diffusion approximation does not hold. This is a preliminary suggestion, and more work is needed to develop its theoretical basis and empirical quantification.

For the moment, we believe that the simplest way to achieve the goals of MCT – to assess the persistence of species in a way that allows a quantitative comparison between different systems (e.g., forests) and a quantitative assessment of the relative contributions of different stabilising mechanisms (e.g., EC covariance vs. relative nonlinearity) – is by simulation of an appropriately-calibrated model. These simulations can be used to estimate metrics that more directly measure persistence, such as the mean time to absorption, the chance of invasion of a species in a community, and species turnover rates.

## v. ACKNOWLEDGMENTS

We thank Michael Kalyuzhny for many helpful discussions and for a critical reading of this manuscript. This research was supported by the ISF-NRF Singapore joint research program (grant number 2669/17).

## Supplementary information

### I. THE LOTTERY MODEL WITHOUT DEMOGRAPHIC STOCHASTICITY

#### A. The diffusion approximation

In this supplement we review the features of the lottery model presented by Chesson and Warner [18], and analyse the roles of *δ* and *σ*, following the discussion in [22, 29]. Most of the results presented here, and in the next supplement which deals with the effect of demographic stochasticity, are based on the diffusion approximation, so their validity is limited to small-amplitude environmental fluctuations. However, the main qualitative insights appear to be generic.

The lottery model (see main text) satisfies the dynamics

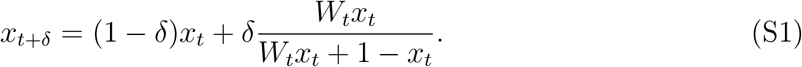

Here we consider the case where the model is symmetric, E[ln *W*_*t*_] = 0, so the mean fitness of the focal species is equal to the mean fitness of its rival. We parameterise 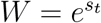, with *s*_*t*_ being any random variable with mean zero and standard deviation *σ*.

If *σ*^2^ ≪ 1, so that the diffusion approximation holds, Eq. (3) yields

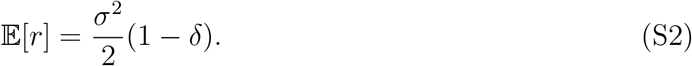

As we shall see below, another important parameter is the strength of the stochastic abundance variations [30]

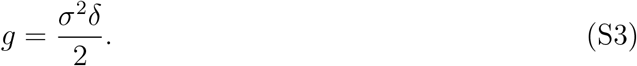

Many of our results turn out to depend on the ratio between these two parameters,

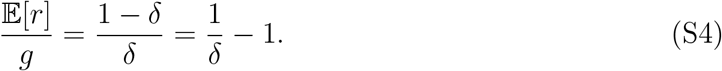

When the diffusion approximation holds, the probability to find the focal species with frequency *x* in the absence of demographic stochasticity follows the Fokker-Planck (forward Kolmogorov) equation [29],

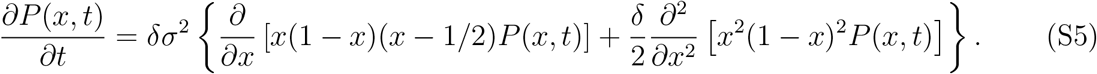

##### 1. Mean time to extinction

If a coexistence state exists, then starting from an arbitrary initial condition, the system eventually relaxes to the steady state *P*_eq_(*x*) for which the time derivative of *P* is zero. The Fokker-Planck equation (S5) then implies

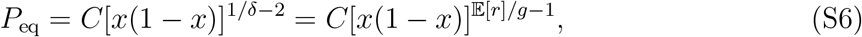

where *C* = Γ(2/*δ* − 2)/Γ^2^(1/*δ* − 1) is the normalisation constant, written in terms of Gamma functions [29].

The rate of extinction events is proportional to the chance to find the system in the extinction zone *x* ≤ *ϵ*, where a natural choice is *ϵ* ≈ 1/*N* [23]. After equilibration,

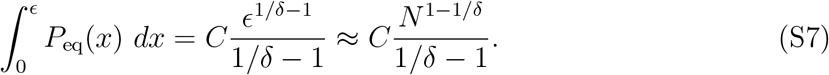

As long as *δ* < 1 this expression goes to zero when *N* → ∞, so the chance to find the system in the extinction zone is arbitrarily small. If *δ* = 1 the equilibrium distribution (S6) does not exist (it is not normalisable). This scenario is discussed in [22], but is less relevant to the cases considered here.

If the mean time to extinction (or the time to absorption) is inversely proportional to the chance to find the system in the extinction zone after equilibration (i.e., if one may neglect the equilibration time), then this time scales like *N*^1/*δ*−1^ = *N*^𝔼[*r*]/*g*^. In that case the time to extinction becomes independent of *σ*, the strength of the environmental variations. Since *g* > 0 by definition, the time to extinction diverges with *N* as long as 𝔼[*r*] > 0, in agreement with [24, 25].

The *σ*-independence of *P*_eq_ is easy to understand from Eq. (S5): one may absorb the overall factor *δσ*^2^ by rescaling the units of time. Accordingly, when the diffusion approximation holds *σ* controls only the equilibration time. Indeed, in panel (b) of Fig. 1 the time to absorption has a simple 1/*σ*^2^ behaviour in the small-*σ* regime.

##### 2. The chance of invasion

The chance of invasion is a time-independent quantity, and in panel (a) of Fig. 1 its dependence on *σ* is shown to be weak and non-monotonic. To understand better this quantity, let us consider first the problem of a particle diffusing along the *y*-axis with velocity 𝔼[*r*] and diffusion constant *g* (this being a standard convection-diffusion problem). The probability *Q*(*y*) to find the particle at *y* satisfies the Fokker-Planck equation

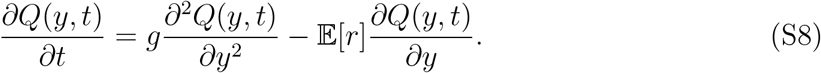

In this case, if the particle starts at *y*_0_, its chance to reach *y* = *L* before it reaches *y* = 0 is given by [see [31], Eq. (2.3.8)]

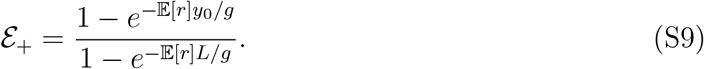

The connection between this result and the chance of invasion becomes clear when one examines the dynamics of the lottery model, Eq. (S5), in the low frequency (*x* ≪ 1) regime,

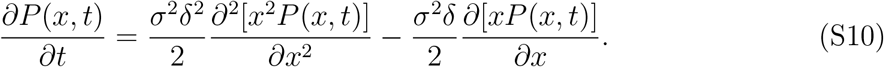

To make Eq. (S10) a simple convection-diffusion equation like (S8), let us consider its dynamics in the log-frequency domain. We substitute *y* = ln *x* and use *Q*(*y, t*) for the probability distribution function in terms of *y*, so that *P* (*x, t*) = *Q*(*y, t*)/*x*, to ensure *P* (*x, t*)*dx* = *Q*(*y, t*)*dy*. We also switch to measuring time in units of the generation time (such that *Q*(*y, t* + *δ*) *≈ Q*(*y, t*) + *δ∂Q*(*y, t*)/*∂t*). This yields

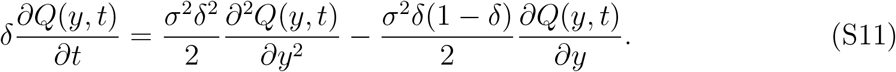

Eq. (S11) has the same structure as Eq. (S8). In an invasion experiment, the extinction point is at *x* = *ϵ*, the initial frequency is *x*_0_ = 2*ϵ*, and successful invasion is declared when the frequency reaches *x*_1_ before it reaches *ϵ*. The corresponding log-frequency values are *y* = ln *ϵ* for extinction, *y*_0_ = ln 2 + ln *ϵ* and *L* = ln *x*_1_. Since the problem is translation-invariant, we can shift the *y* values by ln *ϵ*, so that extinction takes place at zero, *y*_0_ = ln 2, and *L* = ln(*x*_1_/E). Plugging these parameters into Eq. (S9) yields Eq. (6) of the main text. This prediction agrees very well with the measured chance of extinction in Fig. 1 of the main text, and, as one sees, is independent of *σ*. As *σ* grows the diffusion approximation becomes invalid and some weak dependence on *σ* appears.

#### B. Discontinuities in the chance of invasion – Fig. 1(a)

The origin of the discontinuities in the chance of invasion [Fig. 1, panel (a)] may be traced to the chance of extinction in a single step or in a few steps. For example, if *x*_*t*+1_ is given by (2) with *W*_*t*_ = *e^±σ^* and *x*_*t*_ = 2*ϵ*, then as long as

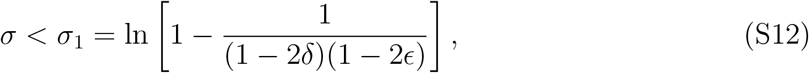

the system cannot reach extinction (*x* = *ϵ*) in a single step. When *δ* < 1/2 there is no solution for *σ*_1_, but for higher values there is a critical value of *σ* above which extinction may take place in a single step, and at this value one observes a sharp decrease in the chance of invasion. A similar argument works for the chance to go extinct in two steps, three steps and so on. For example, for *δ* = 0.7 and *ϵ* = 0.001, one expects a jump at *σ* = 1.254 which corresponds to 𝔼[*r*] = 0.228 [Eq. (3)], precisely as seen in Fig. 1(a).

### II. THE LOTTERY MODEL WITH DEMOGRAPHIC STOCHASTICITY

#### A. Numerical procedure

We would like to introduce a model with demographic stochasticity which corresponds to the lottery model, i.e., a model with a community size of *N* individuals which (for any finite frequency *x*) converges to the two-species equation (2) in the limit *N* → ∞.

At the beginning of each time-step the abundance of species 1 is *n* (its frequency being *x* = *n/N*) and the abundance of species 2 is *N − n* (with its frequency being 1 − *x*). In each time-step the death toll of species 1 is drawn at random from a binomial distribution in which the number of trials is *n* and the chance of success is *δ*, denoted by *B*_*n*_(*δ*), while for species 2 the same holds with *N* − *n* replacing *n* [so that the number of deaths is *B*_N−*n*_(*δ*)].

If *d*_1_ and *d*_2_ denote the death tolls for species 1 and 2, respectively, in a given time step, then the total number of subsequent births (or recruitment events) is *d*_1_ + *d*_2_, to preserve the total population of the entire two-species community. Of these births, the share of species 1 is taken randomly from a binomial distribution wherein the total number of trials is *d*_1_ + *d*_2_ and the chance of success is determined by the relative fitness and abundance of species 1, i.e., the number of new recruits for species 1 is 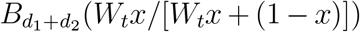. Species 2 wins all the remaining slots.

If the outcome of each binomial trial is replaced by its mean, the dynamics converge to those of Eq. (2) as *N → ∞*. The standard deviation of the relevant binomial trials leads to demographically-induced fluctuations which scale like 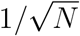, so in the large-*N* limit the two models coincide. However, for small *N*, the effect of demographic stochasticity becomes important (see the next subsection). Moreover, as discussed in the main text, the results of invasion experiments in which the initial condition is *x* = 1/*N* never converge to the results of Eq. (2).

#### B. The diffusion approximation and the relationships between *σ* and *N*

Under pure demographic stochasticity, the Fokker-Planck equation is [21]

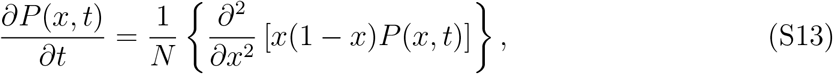

so the equation for a system with both demographic and environmental stochasticity is

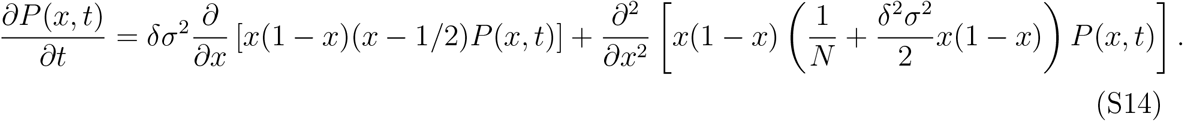

To derive this formula directly one may use the same analysis that leads to Eq. (7) of [32], when the parameter *θ* that was used there to describe mutations vanishes.

Formally speaking, Eq. (S14) does not admit a stable probability distribution function *P*_eq_(*x*), as under demographic stochasticity the system reaches eventually either fixation or extinction. Nevertheless, if the rate of extinction is small enough, the system reaches a quasi-stationary probability distribution which “leaks” very slowly towards extinction or fixation and satisfies *dP* (*x*)/*dt ≪* 1 (see [33] for a formal discussion of this issue).

When the 1/*N* term in Eq. (S14) is negligible (large-*N* limit), the dynamics reduce back to Eq. (S5), *σ* factors out, and the steady state is *σ*-independent. The quasi-stationary probability distribution function becomes *σ*-dependent only when the 1/*N* term is not negligible with respect to *σ*^2^*δ*^2^.

In Eq. (S5) the ratio between the deterministic growth term (the first derivative with respect to *x*) and the random diffusion term (the second derivative) is 1/*δ*, so the growth becomes more and more pronounced as *δ →* 0, and the chance of invasion grows as *δ* decreases and 𝔼[*r*] increases. Here, on the other hand, the amplitude of diffusion is bounded from below by 1/*N*, so as *δ* approaches zero the relative importance of the growth term and the chance of invasion decrease, despite the increase in the value of 𝔼[*r*]. This phenomenon is demonstrated in the inset of Fig. 2(a).

In panel (b) of Fig. 2 of the main text, one observes a strong effect of *σ* on the time to absorption in the small-*σ* regime, followed by saturation as *σ* grows. The same behaviour has been observed for the chance of invasion: while in panel (a) of Fig. 2 we present only the small-*σ* regime for fixed *δ*, we find (results not shown) that the *σ*-dependence disappears when *σ* becomes large. These phenomena are explained by Eq. (S14): when *σ* is large enough, the 1/*N* factor in the diffusion term is negligible, (S14) reduces to (S5) and *σ* factors out.

### III. THE Δ*I*_*b,i*_ METRIC

Ellner et al. [11] have used the original lottery model of Chesson and Warner [18], where the dynamics for one time step (say, a year) are given by

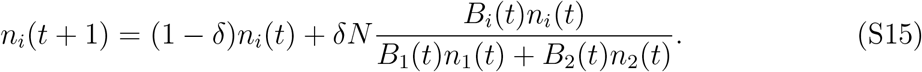

Here *n*_*i*_(*t*) and *B*_*i*_(*t*) are the population abundance and the fitness (given by the fecundity) of species *i* in year *t*, respectively, *δ* is the fraction of the individuals in the community dying in each year, and *N* equals *n*_1_ + *n*_2_. *B*_*i*_(*t*) is picked independently every year from a certain distribution, so one year is the correlation time of the environmental conditions.

The technique of Ellner et al. is based on measuring the growth rates for each species twice, with and without covariance between the environment and the strength of competition (EC covariance). For this purpose, the environment *E*_*i*_ of species *i* is identified with its fecundity *B*_*i*_(*t*), and its competition strength *C*_*i*_(*t*) is defined as [*B*_1_(*t*)*n*_1_(*t*)+*B*_2_(*t*)*n*_2_(*t*)]/(*δN*) [so that *C*_1_(*t*) = *C*_2_(*t*)]. The (logarithmic) population growth rate *r_i_*(*t*) of species *i* is then defined as

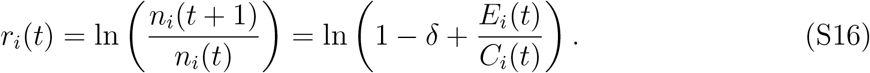

In the case when species 1 invades species 2, for instance, we have *n*_1_(*t*) ≪ *n*_2_(*t*) ≃ *N*, so that *C*_1_(*t*) = *C*_2_(*t*) ≃ *B*_2_(*t*)/*δ*, and Eq. (S16) becomes

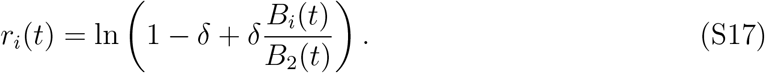

The authors identify the mean growth rate when rare, i.e., 𝔼[*r_i_*(*t*)], with invasibility. Their goal is to measure how much of the invasibility is caused by EC covariance (note that there is an overall 1/*δ* factor between Eq. (S17) and Eq. (3), but it has no effect on the analysis).

Once the simulations are run according to the lottery model and the above calculations performed, the *r_i_*(*t*)’s found from Eq. (S17) are the growth rates *with* the competition strength varying with the environment. To find the corresponding growth rates *without* such EC covariance, the same process is carried out, except with the environment variables given new random values – drawn from the same distribution as before – but with the values of the competition strength fixed to the same values as before. With the new values marked with a ‘#’ superscript, such that 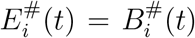, the new growth rates in the case when species 1 invades species 2 become

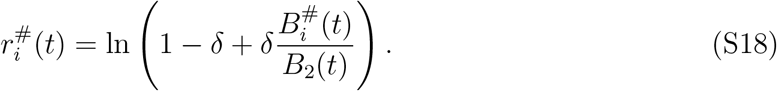

Note that while *B_1_(t)* and *B*_2_(*t*) in Eq. (S17) may be correlated, 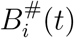 and *B_2_(t)* in Eq. (S18) are not. So the mean growth rate 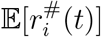 is assumed to correspond to the invasibility when all the parameters are kept fixed except for the removal of EC covariance.

Finally, the contribution of EC covariance to the invasibility, which is a quantitative measure of the storage effect, is defined as

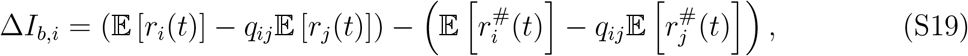

where *i* = 1, 2, *j* ≠ *i* and *q_ij_* are, in general, scaling factors, which for the symmetric lottery model described above equal 1. To find the community-wide counterpart of this measure, one has to sum up Δ*I*_*b,i*_ for the cases when each species is an invader and the others are residents, that is, have much larger initial populations.

### IV. THE *A*_*ij*_*A*_*ji*_ METRIC AND ITS APPLICATION TO A FOREST MODEL

Usinowicz et al. [17, 19] described a forest model in which the lifetime of each tree is divided into two effective stages, a sapling stage and an adult stage. The sapling density *s*_*i*_(*t* + 1) of species *i* at time *t* + 1 (with time measured in years) is given by,

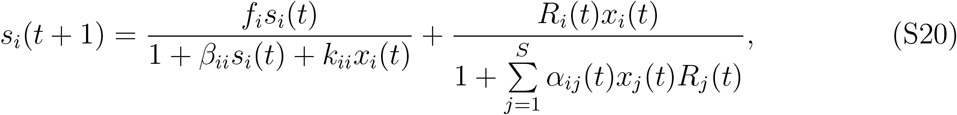

where *x*_i_(*t*) is the adult frequency of species *i* at time *t* and *S* is the total number of species. The first and second terms on the right hand side correspond to the contribution to the sapling population from the surviving saplings from the previous years and from the new saplings generated by the existing adults of the species, respectively. Here *f*_*i*_ is the fraction of the saplings of species *i* surviving from one year to the next, and *R*_*i*_(*t*) is the time-dependent rate at which the adults of species *i* generate saplings. The coefficients *β*_*ii*_, *k*_*ii*_ and *α*_*ij*_ quantify the competition to the saplings of species *i* from the saplings or adults of species *i* or from the saplings of the other species. Hereon, like in [17, 19], the values of these parameters are taken to be *α*_*ij*_ = 1 and *β*_*ii*_ = *k*_*ii*_ = 0 for all *i, j*, in order to isolate the effect of the environmental variations.

The other stage, that of adulthood, is modelled as a process similar to the lottery model [18],

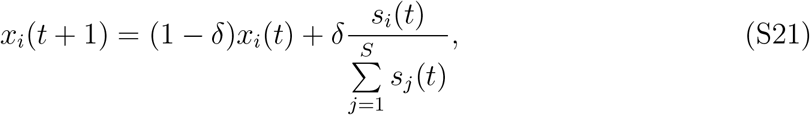

where *δ* is the fraction of the adult population dying at each time-step and the chance of each species to fill an open gap in the adult population is proportional to the relative frequency of its saplings. When *f_i_* = 0 for every *i* this model converges to the lottery model of Eq. (2), so the main difference between the forest and the lottery dynamics is the long-term “memory” associated with sapling survival.

With this setup, the primary measure of coexistence, when the model is limited to competition between only two species *i* and *j*, was suggested to be *A*_*ij*_*A*_*ji*_, where each factor in this product is given by,

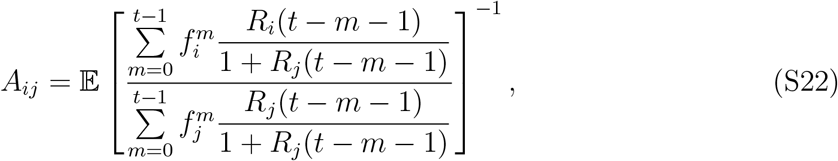

where 𝔼 denotes the expectation value. (Note that in [17, 19] there is a typographical error, such that the sums in the above expression are stated incorrectly to run from *m* = 1 to *t*, instead of from *m* = 0 to *t* − 1.) Unlike Δ*I*_*b,i*_, *A*_*ij*_*A*_*ji*_ is held to be inversely proportional to persistence: the smaller the value of *A*_*ij*_*A*_*ji*_, the more persistent is the system.

To analyze the effect of changing the amplitude of the environmental stochasticity while keeping all other properties of the dynamics intact, we transform the sapling recruitment rates *R*_*i*_(*t*) in Eq. (S20) as

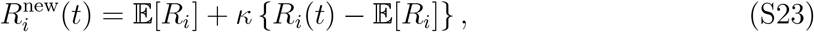

where 𝔼[*R*_*i*_] is the mean value of *R_i_*(*t*) over all years. For *κ* = 0 there is no environmental stochasticity, 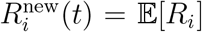, while *κ* = 1 corresponds to the environmental stochasticity in the original dataset, 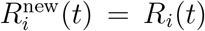. The above transformation preserves the means and correlations amongst all *R*_*i*_(*t*)’s while modulating their amplitudes *σ*. For large enough *κ*-values, this results in some 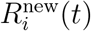 becoming negative (when *R*_*i*_(*t*) < 𝔼[*R_i_*]). For the recruitment rates for *Spondias mombin* and *Spondias radlkoferi* as obtained from [19], this happens at *κ* ≥ 1.171, so we restrict our plots in Fig. 3 to smaller *κ*-values.

The definition of (S22) is presented in [19] (Eq. (D-4) in appendix D) and was obtained through a small-*δ* expansion of the logarithmic growth rate. In that work *δ* was taken to be very small (*δ* = 0.025). As a result, the times to extinction were so long, and the chance of invasion so close to one, that we failed to obtain a reliable estimate of these metrics from numerical experiments. To overcome this difficulty, we used, for Fig. 3, *δ ≥* 0.15. In this case the small-*δ* approximation is problematic, and we have therefore also calculated the value of *A_ij_* using the original definition of 𝔼[*r*] given in Eq. (D-1) of [19]. The differences between the values of *A_ij_* obtained directly using Eq. (D-1) and those obtained from the approximate expression (S22) were tiny in the parameter regime we have considered. In Fig. 3 we present the values of *A*_*ij*_*A*_*ji*_ as obtained from Eq. (S22).

## References

[1] G. E. Hutchinson, The American Naturalist 95, 137 (1961).

[2] D. Tilman, Resource competition and community structure, 17 (Princeton university press, 1982).

[3] R. M. May, Nature 238, 413 (1972).

[4] P. Chesson, Annual Review of Ecology and Systematics 31, 343 (2000).

[5] P. Chesson, Theoretical Population Biology 45, 227 (1994).

[6] P. Chesson, Theoretical Population Biology 64, 345 (2003).

[7] G. Barabás, R. D’Andrea, and S. M. Stump, Ecological Monographs 88, 277 (2018).

[8] S. P. Ellner, R. E. Snyder, P. B. Adler, and G. Hooker, Ecology Letters 22, 3 (2019).

[9] P. Chesson, Journal of Ecology 106, 1773 (2018).

[10] P. L. Chesson, Philosophical Transactions of the Royal Society of London. Series B: Biological Sciences 330, 165 (1990).

[11] S. P. Ellner, R. E. Snyder, and P. B. Adler, Ecology Letters 19, 1333 (2016).

[12] R. A. Chisholm, R. Condit, K. A. Rahman, P. J. Baker, S. Bunyavejchewin, Y.-Y. Chen, G. Chuyong, H. Dattaraja, S. Davies, C. E. Ewango, et al., Ecology Letters 17, 855 (2014).

[13] M. Kalyuzhny, E. Seri, R. Chocron, C. H. Flather, R. Kadmon, and N. M. Shnerb, The American Naturalist 184, 439 (2014).

[14] M. Kalyuzhny, R. Kadmon, and N. M. Shnerb, Ecology Letters 18, 572 (2015).

[15] T. Fung, J. P. O’Dwyer, K. A. Rahman, C. D. Fletcher, and R. A. Chisholm, Ecology 97, 1207 (2016).

[16] A. D. Letten, M. K. Dhami, P.-J. Ke, and T. Fukami, Proceedings of the National Academy of Sciences 115, 6745 (2018).

[17] J. Usinowicz, C.-H. Chang-Yang, Y.-Y. Chen, J. S. Clark, C. Fletcher, N. C. Garwood, Z. Hao, J. Johnstone, Y. Lin, M. R. Metz, et al., Nature 550, 105 (2017).

[18] P. L. Chesson and R. R. Warner, The American Naturalist 117, 923 (1981).

[19] J. Usinowicz, S. J. Wright, and A. R. Ives, Ecology 93, 2073 (2012).

[20] R. Lande, S. Engen, and B.-E. Saether, Stochastic population dynamics in ecology and conservation (Oxford University Press, 2003).

[21] S. Karlin and H. E. Taylor, A second course in stochastic processes (Elsevier, 1981).

[22] A. Dean and N. M. Shnerb, BioRxiv p. 725341 (2019).

[23] P. L. Chesson, Journal of Mathematical Biology 15, 1 (1982).

[24] S. J. Schreiber, M. Benaïm, and K. A. Atchadé, Journal of Mathematical Biology 62, 655 (2011).

[25] S. J. Schreiber, Journal of Difference Equations and Applications 18, 1381 (2012).

[26] T. Spanio, J. Hidalgo, and M. A. Munñoz, Physical Review E 96, 042301 (2017).

[27] A. H. O. Wada, M. Small, and T. Vojta, Physical Review E 98, 022112 (2018).

[28] Y. Yahalom and N. M. Shnerb, Physical Review Letters 122, 108102 (2019).

[29] M. Danino, N. M. Shnerb, S. Azaele, W. E. Kunin, and D. A. Kessler, Journal of Theoretical Biology 409, 155 (2016).

[30] I. Meyer and N. M. Shnerb, Scientific Reports 8, 9726 (2018).

[31] S. Redner, A guide to first-passage processes (Cambridge University Press, 2001).

[32] M. Danino and N. M. Shnerb, Physical Review E 97, 042406 (2018).

[33] D. A. Kessler and N. M. Shnerb, Journal of Statistical Physics 127, 861 (2007).

